# Sulfonylurea receptor coupled conductances alter the performace of two central pattern generating circuits in *Cancer borealis*

**DOI:** 10.1101/2024.07.09.602760

**Authors:** Sonal Kedia, Naziru M. Awal, Jackie Seddon, Eve Marder

## Abstract

Neuronal activity and energy supply must maintain a fine balance for neuronal fitness. Various channels of communication between the two could impact network output in different ways. Sulfonylurea receptors (SURs) are a modification of ATP-binding cassette proteins (ABCs) that confer ATP-dependent gating on their associated ion channels. They are widely expressed and link metabolic states directly to neuronal activity. The role they play varies in different circuits, both enabling bursting and inhibiting activity in pathological conditions. The crab, Cancer borealis, has central patterns generators (CPGs) that fire in rhythmic bursts nearly constantly and it is unknown how energy availability influences these networks. The pyloric network of the stomatogastric ganglion (STG) and cardiac ganglion (GC) control rhythmic contractions of the foregut and heart respectively. Pharmacological manipulation of SURs results in opposite effects in the two CPGs. Neuronal firing completely stops in the STG when SUR-associated channels are open, and firing increases when the channels are closed. This results from a decrease in the excitability of pyloric dilator (PD) neurons, which are a part of the pacemaker kernel. The neurons of the CG, paradoxically, increase firing within bursts when SUR-associated channels are opened, and bursting slows when SUR-associated channels are closed. The channel permeability and sensitivities analyses present novel SUR-conductance biophysics, which nevertheless change activity in ways reminiscent of the predominantly studied mammalian receptor/channels. We suggest that SUR-associated conductances allow different neurons to respond to energy states in different ways through a common mechanism.

## Introduction

Neuronal activity is energetically demanding, and ATP is crucial for maintaining membrane potentials and neurotransmitter pools^1^. It follows that activity rates must need to scale with ATP availability to sustain firing and neuronal health^2,3^. ATP levels and turnover rates could thus be a readout of activity rates and form a feedback mechanism for neurons to sense their own firing level deviations. Neuronal metabolism has been found to play a role in measuring and regulating firing rate homeostasis^4,5^. Ion channel modifications that allow channel gating by ATP/ADP levels are a way to connect metabolic state to activity directly and can have important implications for controlling circuit output.

Sulfonylurea receptors (SURs) are a unique form of the superfamily of ATP binding cassette (ABC) transporter proteins that impart associated ion channel subunits with gating dependent on intracellular concentrations of ATP ([ATP])^6^. The two SUR isoforms are widely expressed in mammalian tissues such as the brain, heart and skeletal muscle^7^. ATP-sensitive K^+^ channels (K-ATP), first discovered in the pancreas, are complexes of SURs and inward-rectifying K^+^-channel (K_IR_) subunits and are the most prevalent SUR-associated channels. They are predominantly closed under conditions of adequate ATP, allowing neuronal firing, and open when these levels fall-hyperpolarizing the membrane potential and preventing further activity^8,9^. Thus, they provide an important way to link a cell’s metabolic state to its activity. Associations are also reported between the pore-forming subunit of the calcium-activated transient receptor potential melastatin 4 (TRPM4) channel and SURs forming functional channels which are cation non-selective and lead to depolarization of neuronal membranes and a large influx of ions resulting in osmotic swelling^10^. The two forms play different roles in the nervous system. K-ATP channels have been studied in the context of preventing epileptic activity^11^, enabling burst firing in dopaminergic neurons^12^, and allowing glucose-sensing by hypothalamic neurons^13^; and are largely considered protective^14^. TRPM4-SURs are implicated in neuronal damage and edema following various forms of injury^15^.

[ATP]_i_ responsiveness in nervous systems must have some universal features across evolution. Consequences of SUR activity as a mechanism for ATP responsiveness have largely been studied in vertebrate systems; and it is unclear how this same problem is solved in earlier animals and as a function of circuit architecture. Invertebrates are ectothermic and their metabolic rates have temperature dependencies which could generate additional challenges for energy-regulated feedback mechanisms in their nervous systems.

We tested the action of pharmacological agents that bind to SURs on spontaneous bursting activity in two central pattern generators (CPGs) in the crab, Cancer borealis, which allows us to probe energy-related firing dynamics in two spontaneously active networks. CPGs have unique advantages for the exploration of circuit design and regulation, key amongst which are their easily evoked patterned rhythmic output and the direct relationship between this output and the final motor behavior elicited^16^. The stomatogastric ganglion (STG) controls rhythmic contractions of the pylorus and stomach teeth in the foregut and the cardiac ganglion (CG) controls heart contractions. Both small networks are remarkably robust while responding to changes in the environment. Pharmacological agents that bind to SURs affect opening and closing of the associated channels were discovered well before the proteins or channels were identified and have been well characterized and used extensively in clinical settings ^17,18^. We used these agents to explore the presence and role of SUR-associated channels in the two crab CPGs.

## Materials and methods

### Experimental animal and preparation details

Adult male Jonah crabs (Cancer borealis) were purchased from Commercial Lobster (Boston, MA) and held without food in artificial seawater at 10–13°C on a 12 h light/12 h dark cycle. Animals were held on ice for 30 min prior to dissection. Independent dissections were performed for the CG and STG as previously described. For the STG, the stomach was dissected from the animal. The intact stomatogastric nervous system (STNS) was isolated from the stomach, including: the two bilateral commissural ganglia, esophageal ganglion, and STG, with connecting motor nerves. The STNS was pinned down in a Sylgard-coated Petri dish (10 ml) and continuously superfused with chilled saline (440 mM NaCl, 11 mM KCl, 26 mM MgCl_2_, 13 mM CaCl_2_, 11 mM Trizma base, 5 mM maleic acid, pH 7.45 measured at room temperature∼23°C). For the 0.5x [K^+^] experiments, KCl concentration was adjusted to 5.5mM and all other components were the same. For low calcium experiments, the CaCl_2_ concentration was 1.3mM (lower than this impacts the health of the preparation) and the remaining divalent cation concentration was made up by the addition of 11.7mM MnCl_2_. For the CG, the heart was dissected from the animals and the intact cardiac ganglion was isolated from it and similarly pinned in a dish and superfused with saline.

### Electrophysiology

The STG was desheathed and intracellular recordings from somata were performed with 10–30 MΩ glass microelectrodes filled with internal solution (10 mM MgCl2, 400 mM potassium gluconate, 10 mM HEPES buffer, 15 mM NaSO4, 20 mM NaCl^19^ or 0.4M K SO, 3.5mM NaCl, and 25mM KCl for voltage clamp recordings). Intracellular signals were amplified with an Axoclamp 900A amplifier (Molecular Devices, San Jose, CA). Neuronal identity was established after impaling the somata with sharp electrodes based on spiking activity observed on their respective nerves. Extracellular nerve recordings from the CG and STG were made by building wells around nerves using a mixture of Vaseline and mineral oil and placing stainless steel pin electrodes within the wells to monitor spiking activity. Extracellular nerve recordings were amplified using model 3500 extracellular amplifiers (A-M Systems). Data were acquired using a Digidata 1440 digitizer (Molecular Devices, San Jose, CA) and pClamp data acquisition software (Molecular Devices, San Jose, CA; version 10.5). Extracellular nerve recordings were amplified using model 3500 extracellular amplifiers (A-M Systems). Data were acquired using a Digidata 1440 digitizer (Axon Instruments) and pClamp data acquisition software (Axon Instruments, version 10.7). Removal of neuromodulatory inputs to the STG (decentralization), was performed by adding isotonic sucrose (750 mm) and 0.1μm tetrodotoxin (TTX) to a Vaseline well built around the stomatogastric nerve. Current injection ramps were performed in the presence of 10^−5^M picrotoxin (PTX) superfused continuously over the network and voltage clamps recordings used both PTX and TTX in the superfused saline. For current injection ramps and voltage clamp recordings, PD neurons were impaled with two electrodes, one of which was used for current injections and the second for reporting membrane potential.

### Pharmacology

Diazoxide, tolbutamide and glibenclamide were purchased from Sigma-Aldritch (D9035, T0891 and G0639). Stocks were made in DMSO and diluted in saline to the appropriate concentrations for use. Preparations were superfused with the drugs for a minimum of 15 minutes before performing the current injection and voltage clamp experiments.

### Electrophysiology and statistical analyses

Recordings were acquired using Clampex software (pClamp Suite by Molecular Devices, version 10.7) and analyzed with custom MATLAB scripts to calculate burst properties. Traces were low-pass filtered to detect slow wave membrane oscillations. Minimum membrane potentials were measured at the trough of the oscillation. High-pass filters were applied to identify action potentials.

Spike and oscillation thresholds: To calculate spike threshold, spike onset was defined as the voltage corresponding to the maximum curvature of the first derivative of the voltage (dV/dt), when dV/dt crossed the threshold value of 10 mV/ms. Oscillation threshold was identified as the trough value of the first negative peak in the low-pass filtered traces.

Channel properties in voltage-clamp: These were determined through applying voltage steps in two-electrode voltage clamp mode in the different drugs. Currents were measured at the last second of the voltage step, at steady state. The expected E_reversal_ shift for a pure K+ conductance was calculated from the Nernst equation: E_rev_= RT/zF x ln([ion]_out_/[ion]_in_). 5 minute averages were used for data representation. SPSS was used to run statistical tests on data as reported in the paper. N’s represent individual STNS preparations from individual crabs.

### Sequence identification and RNA expression

There is no well-curated reference genome for C. borealis. Therefore tBLASTn searches were run on the C. borealis transcriptome using bait sequences from Drosophila melanogaster and Daphnia pulex to identify a candidate sequence for C.borealis SUR (GeneBank ID:GEFB01014997). The predicted C. borealis sulfonylurea receptor 1 shares 55.03%, 39.01%, and 34.74% sequence identity with Homarus americanus ABCC-9 like isoform (XP_042213487.1), Daphnia pulex SUR 2B (EFX68442.1) and Drosophila melanogaster SUR (NP_001334732.1) respectively.

Tissue from STGs and CGs was collected after dissection and immediately stored in a cryogenic microcentrifuge tube containing 400 µL lysis buffer (Zymo Research) at −80 °C until RNA extraction. Total RNA was extracted using the Quick-RNA MicroPrep kit (Zymo Research) per the manufacturer’s protocol. Following RNA extraction, individual RNA samples were reverse transcribed into cDNA using Superscript III First-Strand Synthesis System (Thermo Fisher) primed with oligo-dTs as per the manufacturer’s protocol in 20-µL reactions. The following primers were designed based on putative sequences identified from the transcriptome: 18SF 5’-AGGTTATGCGCCTACAATGG-3’, 18sR 5’-GCTGCCTTCCTTAGATGTGG-3’; SURF: 5’-GTGACCTAAGACTTGGATGACC, SURR: 5’-GGAACTCGTTAAACACCCGC-3’; KirF: 5’-GCCCGTCTTAATAGCCAGGA-3’, KirR: 5’-ATGGTTGTGGTCCTGCAGTA-3’; TRPM4F: 5’-GTCAAGACTCACGCCTGACA-3’, TRPM4R: 5’-CCTCGTCGTCTTCATCCTCG-3’. PCR reactions were run with STG and CG cDNA to allow for detection of gene expression. PCR products were run on a 2% agarose gel with ethidium bromide and bands were visualized using Bio-Rad Chemidoc imaging system.

## Results

### STG and CG circuitry

The pyloric network of the STG, and the CG are well studied for rhythmic motor pattern generation. The motor neurons in both networks oscillate constantly and ‘semi-autonomously’, with the help of neuromodulatory inputs, and fire bursts of action potentials that elicit muscle contractions. These neurons continue to produce fictive motor patterns in vitro when isolated from the foregut and heart respectively and these preparations can be used for intracellular and extracellular reporting of circuit activity (Fig 1A, D). The pyloric network of the STG is driven by a rhythmically bursting pacemaker kernel consisting of an interneuron, Anterior Burster (AB), and two motor neurons, Pyloric Dilators (PDs). They rhythmically inhibit one Lateral Pyloric (LP) and several Pyloric (PY) neurons (Fig 1B). The synaptic partners of the pacemaker neurons burst on rebound from inhibition to form a characteristic triphasic rhythm in which the dilators (PDs) and constrictors (LP and PYs) are active in turn (Fig 1C). The CG consists of 9 neurons, 4 small cells (SCs) (pacemaker interneurons) and 5 large cells (LCs) (motor neurons) connected both by electrical and excitatory synapses (Fig 1E). SCs burst and drive bursting in the LCs in tandem and form a characteristic ‘heart beat’ that can be recorded extracellularly from the trunk (Fig 1F).

**Figure 1.**
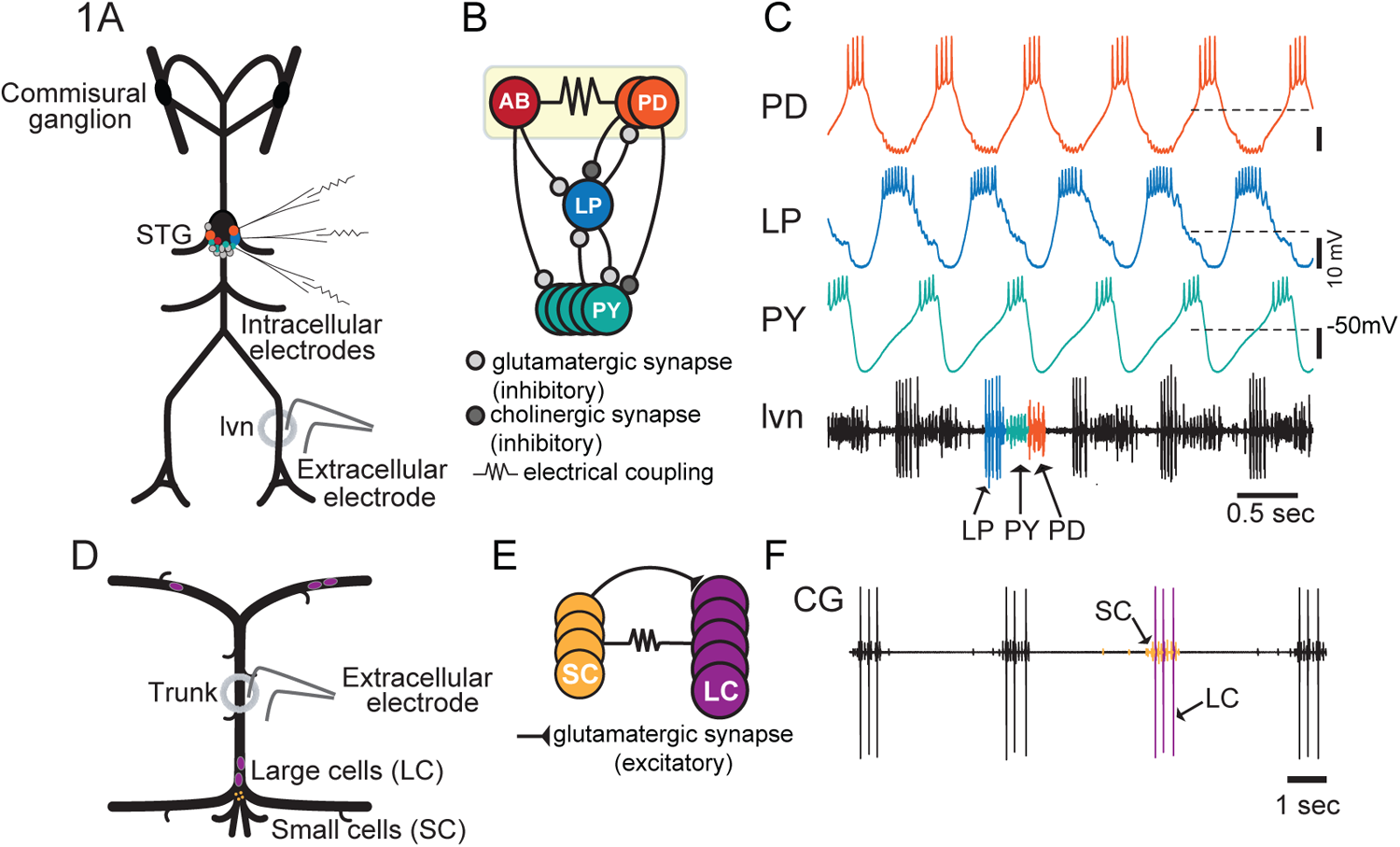
Two C. borealis CPGs. A. Schematic of the experimental setup: in vitro STNS preparation and recording electrodes. B. Wiring diagram of the pyloric network. C. Example voltage traces of intracellular recordings from the three classes of motor neurons of the pyloric circuit and the composite firing sequence of all three recorded extracellularly from the lvn nerve (bottom-most trace). D. Schematic of the CG preparation. E. CG circuit configuration. F. Example extracellular recording of SC and LC burst firing.

### SUR pharmacology

SURs are named for the class of drugs, sulfonylureas, first discovered as hypoglycemic agents and used clinically for years before the K-ATP target was discovered ^17,20^. Numerous sulfonylureas have been developed for their use in diabetes management due to their specificity for SURs through binding sites separate from the nucleotide binding sites^21^. SURs are regulatory subunits and the drugs act as channel closers regardless of the associated channel, mimicking the response to a higher ATP/ADP condition^6^. We studied the response of different concentrations of the sulfonylurea glibenclamide, developed for its high affinity and specificity, on various features of the rhythmic activity of both the pyloric and cardiac networks^12^. We refer to glibenclamide as ‘channel closer’ for ease of communication. A related sulfonamide, diazoxide (referred to as ‘channel opener’), stimulates the opening of known SUR-associated channels^22^. We utilized the existence of this well-established pharmacology to bidirectionally modulate SUR-associated channel activity and study its impact on the stereotyped outputs of the STG and CG.

### Channel opener dose response in the STG

We first tested the channel opener as a means of investigating the presence of SUR-associated channels in STG preparations and the impact of an active SUR-conductance on pyloric output. Increasing concentrations of channel opener, diazoxide, were applied to STG preparations and the responses of PD neurons were recorded (Fig 2A). Burst properties and membrane potentials were analyzed over the time course of drug applications and wash outs from PD recordings (Fig 2B). PD neurons depolarize in the presence of channel opener at all concentrations which initially results in a small increase in burst frequency, as is expected from an appropriately sized depolarizing stimulus. Over the period of drug application however, the effect shifts towards a dramatic reduction in bursting and this effect is more pronounced as the diazoxide concentration increases (Fig 2B). This is consistent with the opening of a large shunting conductance, blocking firing. The channel opener had a dose-dependent effect on PD burst frequencies measured at the final 5 minutes of drug application compared to 5 minutes of baseline recorded immediately before drug application. All preparations burst slower or stopped in 200μM diazoxide, and all stopped bursting entirely in 400μM diazoxide (Fig 2C, Table 1). This corresponded with a decrease in spiking as well (Fig 2D, Table 1) although the membrane potential was depolarized at all concentrations (Fig 2E, Table 1). The effect appears to be network wide; all pyloric neurons stop firing at the highest concentration of the channel opener suggesting that channel expression is widespread in the pyloric network (Fig 2F).

**Figure 2.**
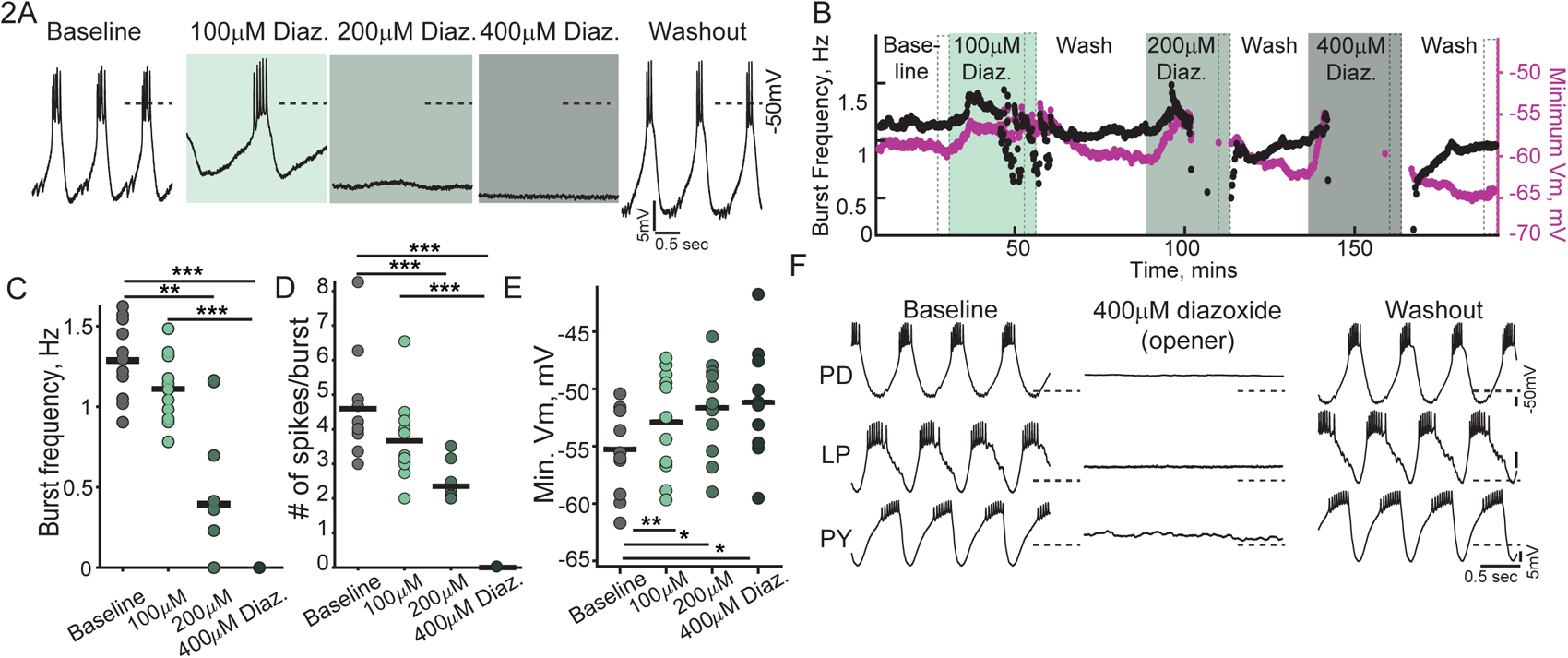
Diazoxide (channel opener) dose response-effects on pyloric circuit activity. A. Example intracellular traces of PD membrane potential in increasing concentrations of diazoxide (diaz.). B. Time course of instantaneous PD burst frequency and minimum membrane potential of oscillation trough over diazoxide applications and wash outs show in panel A. C. Quantification of PD burst frequency. Friedman-Dunn’s test for repeated measures, post-hoc Wilcoxon-signed rank test with Bonferroni’s correction used for C and D. Post-hoc tests-Baseline-200μM: p=0.01; Baseline-400μM: p<0.001, 100-400μM: p=0.01. D. Quantification of # of PD/burst in different diazoxide concentrations. Significant differences between-Baseline-400μM: p<0.001; Baseline-200μM: p=0.001; 200-400μM: p=0.003. E. Quantification of minimum membrane potential of PD compared across groups using a one-way repeated measures ANOVA (F (1.46, 14.66)= 7.01, p=0.011) with post-hoc T-tests with Bonferroni’s correction resulting in significant differences between Baseline-100μM: p<0.006; Baseline-200μM: p=0.018; Baseline-400μM: p=0.028. F. Example traces of all pyloric neurons silenced in the highest concentration of diazoxide. Each dot represents a single preparation. Horizontal bars represent population means. *: p<0.05; **:p<0.01; ***:p<0.001.

**Table 1.** Diazoxide (channel opener) dose response: Means and standard deviations of the burst properties of the pyloric and cardiac circuit 5 minutes before drug application (baseline), 15-20 minutes after diazoxide applications at different concentrations and after washout.

| | Baseline | 100μM<br>diazoxide | 200μM<br>diazoxide | 400μM<br>diazoxide | Washout<br>(>15 mins) |
| --- | --- | --- | --- | --- | --- |
| PD Burst frequency, Hz | 1.29±0.24 | 1.11±0.21 | 0.54±0.61 | 0 | 1.2±0.26 |
| PD # of spikes/burst | 4.6±1.5 | 3.7±1.2 | 1.4±1.2 | 0 | 3.8±1.6 |
| PD Min. membrane V,<br>mV | -55.3±3.8 | -52.9±4.5 | -51.6±4.1 | -51.2±4.7 | -54.6±4.7 |
| CG burst frequency, Hz | 0.16±0.06 | 0.17±0.06 | 0.17±0.06 | 0.2±0.06 | 0.15±0.05 |
| LC # of spikes/burst | 7±2.4 | 9.4±2.7 | 10.6±3.4 | 13.5±6.3 | 8±2.9 |
| SC # of spikes/burst | 31.5±11.2 | 37.9±10 | 42.2±10.1 | 44.9±10.2 | 29.5±8.1 |

### Channel closer dose response in the STG

Pyloric neuron activity in intact preparations was recorded intracellularly in increasing concentrations of the channel closer, glibenclamide, to understand the contribution of open SUR-channels to the baseline rhythm (Fig3A). The channel closer had the opposite effects on pyloric activity as the channel opener. In intact preparations glibenclamide concentrations higher than 100μM produced a modest but significant increase in the burst frequency of the network and in the number of spikes per burst in PD neurons (Fig 3B, C, Table 2). LP neurons did not increase their spiking in the same manner as PD neurons (data not shown). Since the magnitude of change elicited by the closer was small in intact preparations, the effects of the same concentrations of the closer were recorded in pyloric neurons from decentralized preparations (Fig 3D). Decentralization is the removal of neuromodulatory inputs to the STG and this results in a slower and less regular rhythm. Both the burst frequencies and number of spikes per burst in PD neurons increased in higher concentrations of glibenclamide, including an initiation of bursting in non-rhythmic preparations (Fig 3E, F, Table 2). Channel closing produced a much more potent response in decentralized networks than in intact preparations (180% vs 25% increase in burst frequency in 200μM glibenclamide).

**Figure 3.**
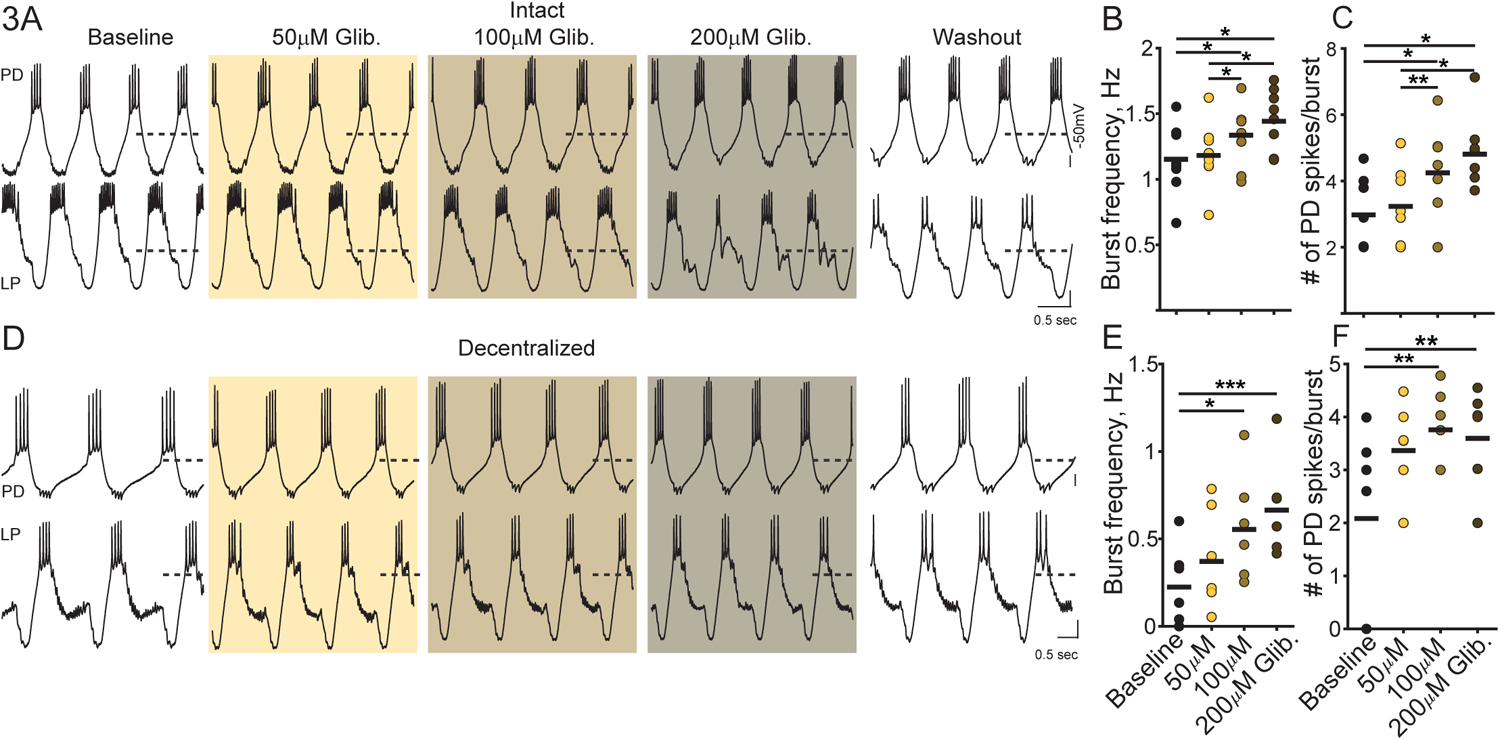
Glibenclamide (channel closer) dose response-effects on pyloric circuit activity. A. Example intracellular traces of PD and LP membrane potential in increasing concentrations of glibenclamide (glib.) in intact preparations. B. Quantification of PD burst frequency in intact preparations compared using repeated measures ANOVA (F(1.49, 11.88) =12.5;p=0.002), post-hoc pairwise T-test with Bonferroni’s correction; Baseline-200μM p=0.027 Baseline-300μM: p=0.036; 100μM-200μM: p=0.019, 100μM-300μM: p=0.028. C. Quantification of # of PD spikes/burst in increasing glibenclamide concentrations using the same statistical tests as B, F(3, 15) =14.95; p<0.001. Significant differences in post-hoc pairwise comparisons between Baseline-200μM glibenclamide: p=0.037; Baseline-300μM glibenclamide: p=0.015; 100μM-200μM glibenclamide: p=0.007; 100μM-300μM glibenclamide: p=0.014. D. Example intracellular traces of PD and LP membrane potential in increasing concentrations of glibenclamide in decentralized preparations. E. Quantification of PD burst frequency in decentralized preparations compared using Friedman-Dunn’s test for repeated measures, post-hoc Wilcoxon-signed rank test with Bonferroni’s correction Baseline-200μM glibenclamide: p=0.022; Baseline-300μM glibenclamide: p=0.001. F. Quantification of # of PD/burst in decentralized preparations in different glibenclamide concentrations, same statistical tests as E. Significant differences between Baseline-200μM glibenclamide: p=0.031; Baseline-300μM glibenclamide: p=0.044. Each dot represents a single, horizontal bars represent population means. *: p<0.05; **:p<0.01; ***:p<0.001.

**Table 2.**
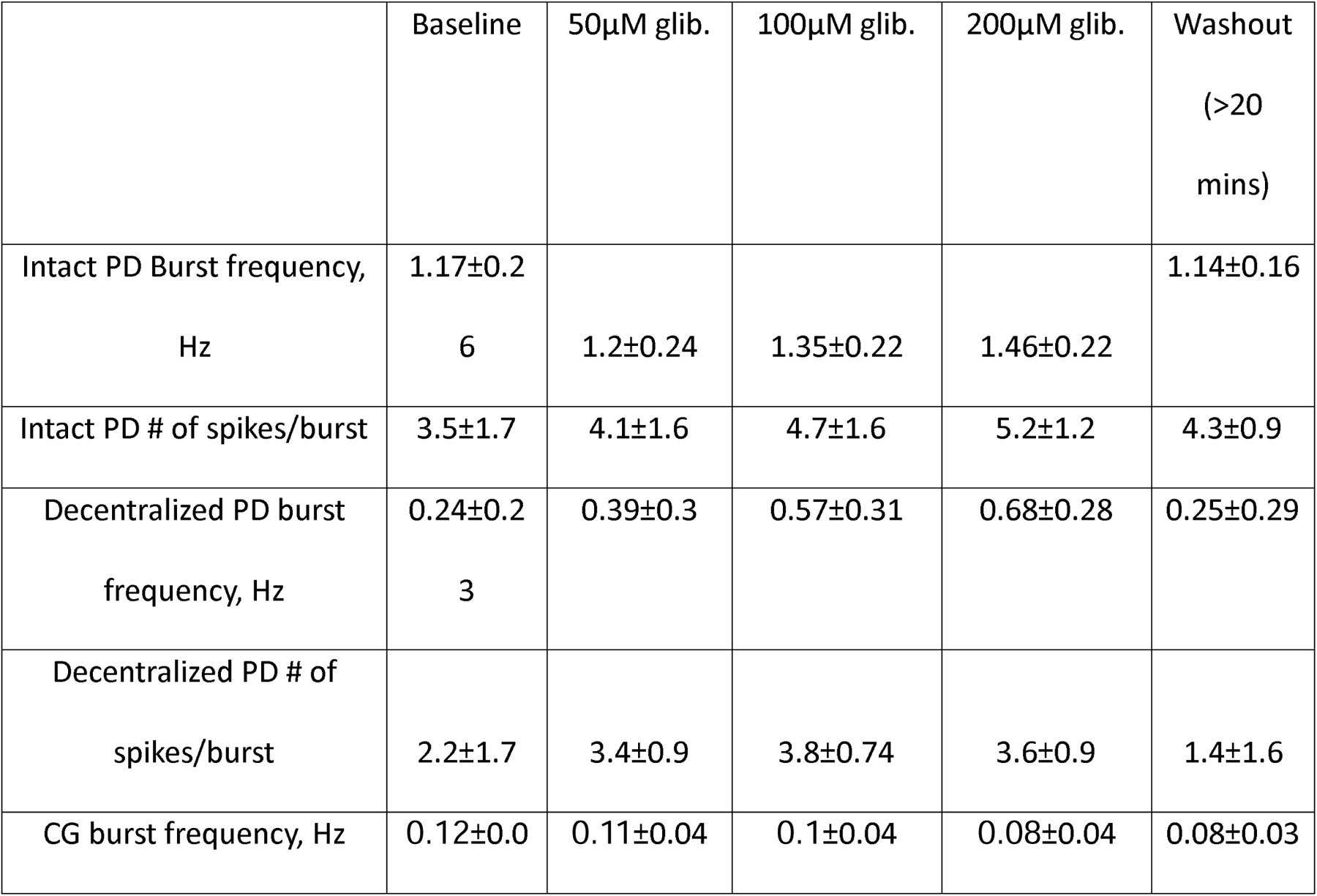
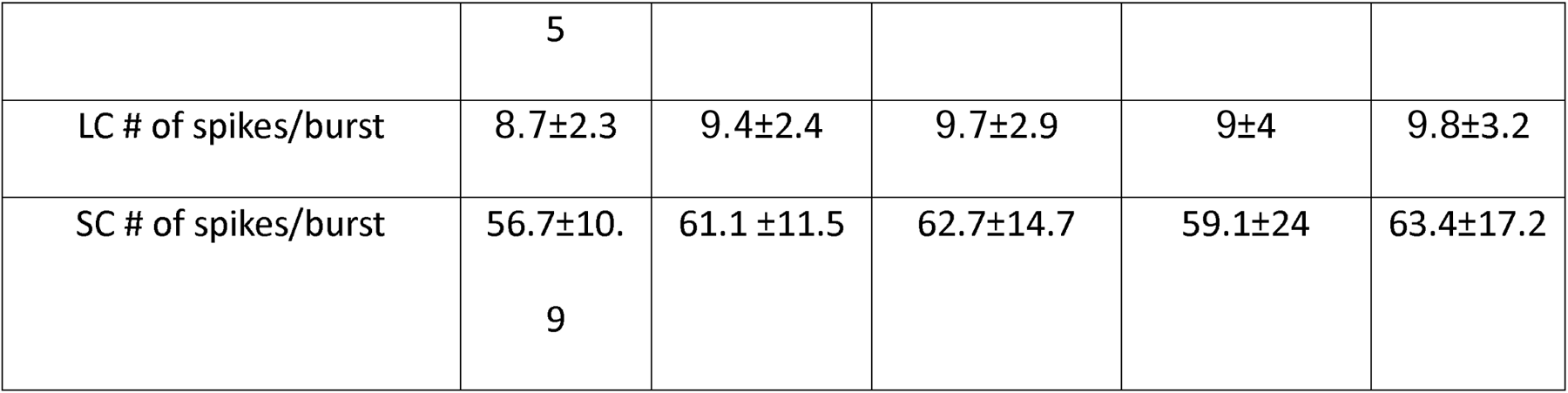
Glibenclamide (channel closer) dose response: Means and standard deviations of the burst properties of the pyloric and cardiac circuit in baseline (before drug application), different concentrations of glibenclamide (glib.) and after washing out glibenclamide.

### Intrinsic properties of PD neurons impacted by SUR-pharmacology

We further examined the influence of channel opening and closing on the intrinsic properties of PD neurons, specifically the spike and oscillation thresholds, to investigate the mechanisms of channel action on network activity. We injected slow ramps of current into PD neurons while recording the membrane potential response in control and then in channel opener (400μM diazoxide) and measured the first instance of spiking and membrane oscillation (Fig 4A). The same protocol was conducted in separate preparations in closer (200μM glibenclamide) (Fig 4B). 2/9 preparations failed to spike with the range of currents we injected in the channel opener and in the remaining 7, spike thresholds moved to significantly depolarized potentials (Fig 4C, left panel); while spike thresholds did not change significantly in the presence of the channel closer (Fig 4C, right panel). 4/9 preparations failed to generate membrane oscillations in the channel opener, even if the PD neuron was firing. The remaining five had a significant depolarized shift in the oscillation threshold in opener compared to baseline (Fig 4D, left panel) and oscillation thresholds were significantly more hyperpolarized in channel closer (Fig 4D, right panel). Therefore, while PD neurons can still spike when the SUR-associated channel is open, it requires a much bigger depolarizing push to make them do so and even then, membrane oscillations may not be restored. Closing the channels on the other hand, allows for membrane oscillations to emerge at more hyperpolarized membrane potentials. These changes were concomitant with changes in input resistance measured at the initial potential drop at the ramp onset. Input resistance decreased in channel opener (Fig 4E, left panel), as would be expected with the opening of a large conductance and consistent with the hypothesis of a shunt. Input resistances increased in channel closer suggesting that some proportion of channels were open in baseline conditions (Fig 4E, right panel).

**Figure 4.**
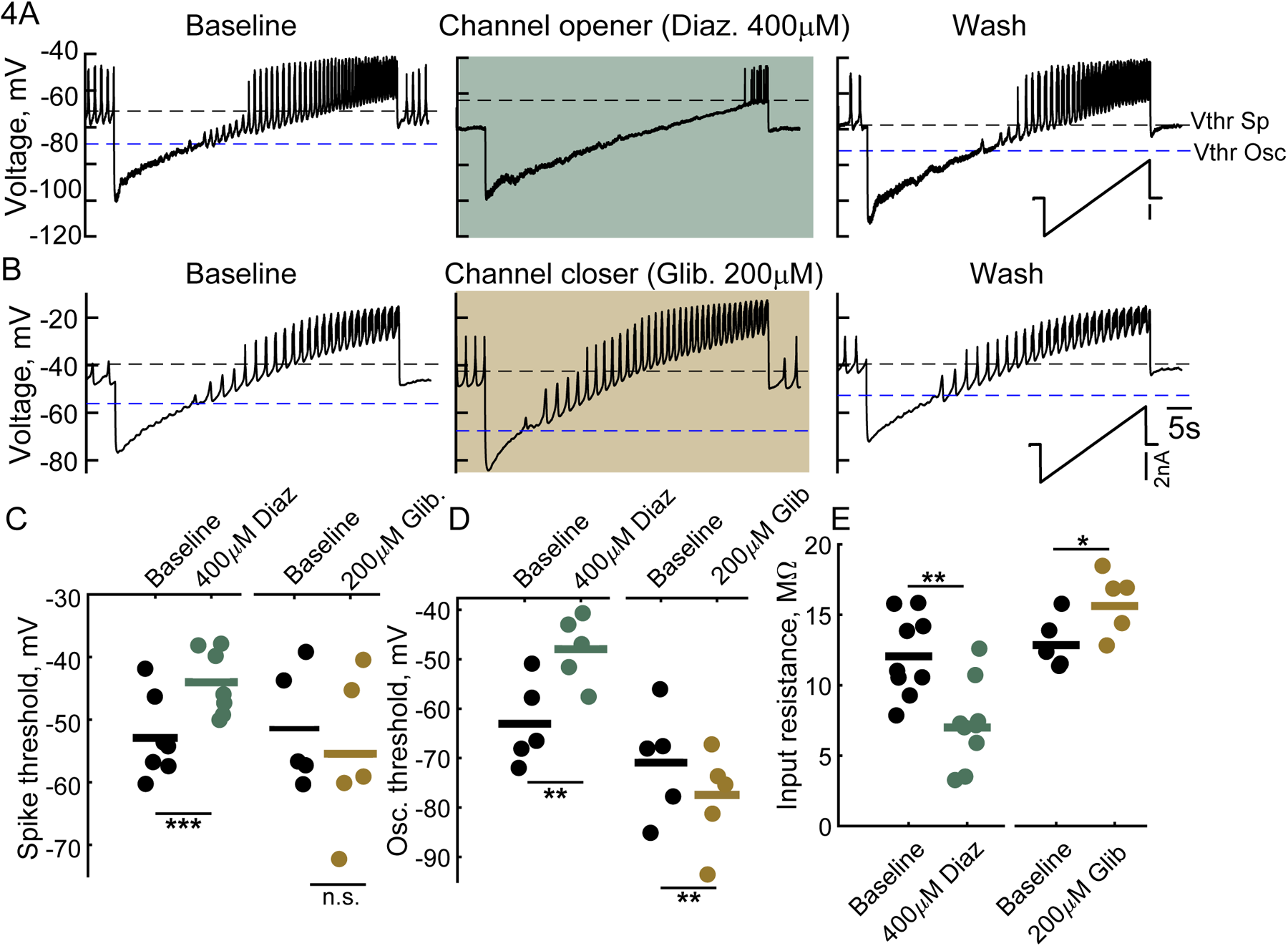
Effects of the SUR-conductance on intrinsic excitability of PD neurons. A, B. Example traces of intracellular PD neuron traces in response to injections of slow current ramps in channel opener (A) and channel closer (B). Current injection protocol is the inset in right panel of B. C. Quantification of spike threshold in baseline(-52.9±6.1mV) and 400μM diazoxide (-44.1±5mV), Paired T-Test p<0.001 (left); and baseline (-51.5±8.4mV) and 200 μM glibenclamide (-55.5±11.4) (right) D. Quantification of oscillation thresholds in baseline (-63.1±7.7mV) and 400μM diazoxide (-48±6.1mV), Paired T-test p=0.012 (left); baseline (-69.6±9.3mV) and 200 μM glibenclamide (76.8±8.8), Paired T-test: p=0.005 (right). E. Quantification of input resistance in baseline (12.1±2.7MΩ) and 400μM diazoxide (7.2±2.8MΩ), Paired T-test p=0.006; and baseline (13±1.7 MΩ) and 200 μM glibenclamide(15.9±2 MΩ), Paired T-test: p=0.031. Each dot represents a single preparation, horizontal bars represent population means. *: p<0.05; **:p<0.01; ***:p<0.001.

### SUR-associated channel properties

The effects of the channel openers necessitated a closer look at channel permeability to establish the identity of the SUR-associated channel. Tolbutamide is a less potent but reversible drug as compared to glibenclamide^21,23^; hence we used tolbutamide for these experiments to ensure a complete wash-out since both drugs were serially applied to the same preparations. We performed two-electrode voltage clamp experiments and gave voltage steps in the presence of the channel opener (400μM diazoxide) and the channel closer (750μM tolbutamide) and measured the injected current at steady state in both cases (Fig 5A, left and middle panel). The difference current obtained from subtracting closed channel traces from open channel traces gives a close approximation of the current passing through the SUR-conductance, as other conductances active at those potentials have been subtracted out (Fig 5A, right panel). IV relationships of the currents from Fig 5A indicate the reversal potential for the difference current obtained (Fig 5B). We recorded an average reversal potential of -65.0±4.7mV from the difference currents measured in 9 preparations (Fig 5C). E_K_ in the STG neurons is reported to be ∼-80mV therefore these results suggest the presence of additional conductances affected by the drugs. To confirm that K^+^ still participates in this conductance, we conducted the same experiments in normal saline and in 0.5x[K^+^] (Fig 5D, E, F). Under these conditions the reversal potential of the difference current shifted by ∼8mV (baseline: -68.2±2.4mV, 0.5x[K^+^] : - 76.5±2.4mV, Paired T-test: p=0.036) (Fig 5G). The Nernst potential for this [K]_o_ change should be ∼17mV shifted for a pure K^+^ conductance. Therefore, we postulate that the SUR-channel in STG neurons is permeable to other ions. TRPM4 channels are calcium sensitive. In some of the same preparations we also conducted the TEVC experiments in low [Ca^2+^] saline (Fig 5D right panel). There was a decrease in the difference current measured under these conditions (Fig 5E) confirming that this conductance too, is calcium sensitive (I in control saline: 7.1+1.5nA; low [Ca^2+^] saline:1.1+1.3nA. Paired T-test: p=0.02).

**Figure 5.**
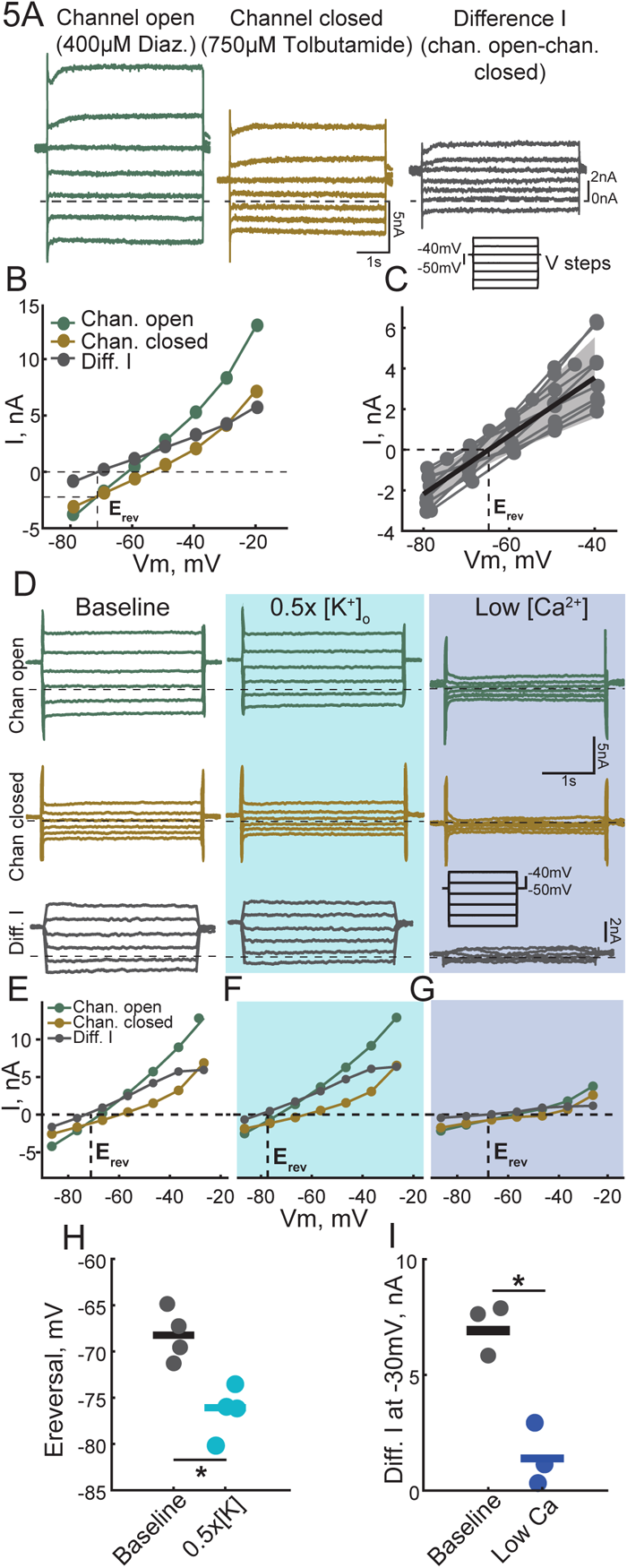
Voltage clamp assessment of channel properties of the SUR-conductance in the STG. A. Example current traces acquired while stepping the membrane potential (inset) in diazoxide (green) and tolbutamide (gold) and the resulting difference current calculated in grey. B. IV plots of the current traces in A with Ereversal (Erev) of the difference I marked. C. IV relationships of the difference current measured across preparations and the average Ereversal measured for the SUR-conductance. D. Example current traces of voltage clamp experiments conducted in diazoxide (green) and tolbutamide (gold) and difference I (grey) in different saline conditions. Left column is control saline recordings. Middle column is recordings made in 0.5x[K^+^] saline and the right column is recordings made in low [Ca^2+^] saline. E. IV plots of left traces in D with baseline Erev. F. IV plots of middle traces in D with Erev in 0.5x[K^+^] . G. IV plots of left traces in D in low [Ca^2+^] . H. Quantification of the Ereversal of the SUR-conductance in control and 0.5x[K^+^] saline across preparations. I. Quantification of the difference I measured at -30mV in control and [Ca^2+^] saline. Each dot corresponds to one preparation, horizontal bars represent population means. *: p<0.05; **:p<0.01; ***:p<0.001.

### SUR pharmacology in the CG

The CG is also a CPG, and it generates rhythmic contractions of the heart muscles, but its wiring scheme differs from the STG (Fig 1B, E). We conducted similar dose responses to both channel opener and closer while recording the CG output extracellularly (Fig 6A, B). Contrary to the results obtained from the STG, channel opening led to an increase in activity in the CG. The largest effect was seen in the number of spikes per burst in both small and large cells, both of which increased in a dose-dependent fashion (Fig 6C, D, Table 1). Burst frequencies were not significantly altered by any concentration of the channel opener (Table 1). The strongest effect of channel closing, however, was a dose-dependent decrease in the burst frequency (Fig 6E, Table 2) whereas the number of spikes per burst did not change significantly in glibenclamide (Table 2). Therefore, while both drugs affected individual neuronal firing and the frequency of bursting in a correlated fashion the STG; these features were decoupled in their response to channel opening and closing in the CG. Channel opening increased spiking within bursts while burst frequencies were not significantly altered while channel closing slowed down the bursts while spiking remained largely unaltered. Channel closing had a more pronounced effect in the CG compared to the STG, suggesting that there may be more open channels contributing to the rhythmic output basally.

**Figure 6.**
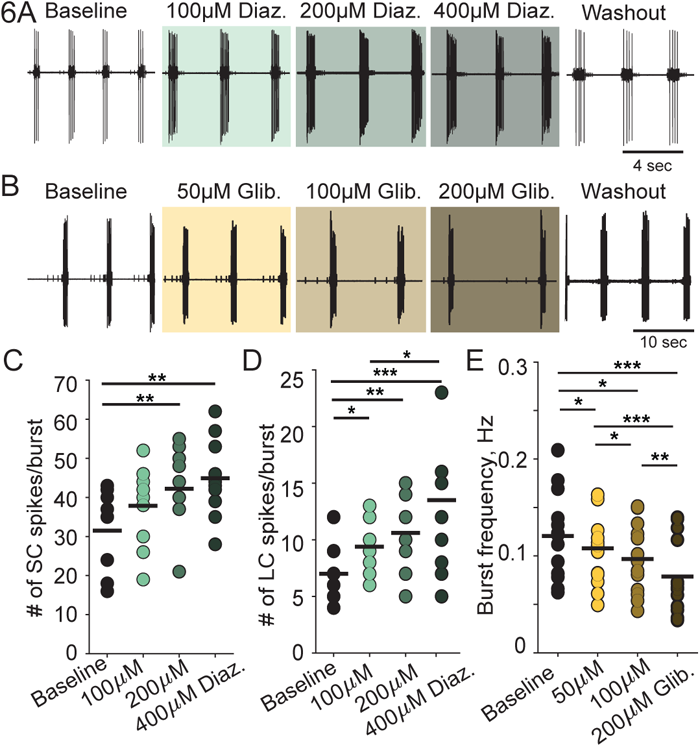
Dose response of diazoxide (channel opener) and glibenclamide (channel closer) on the cardiac ganglion (CG). A. Extracellular recordings from the trunk of the CG in baseline, increasing concentrations of diazoxide (diaz.) and after drug wash out. B. Extracellular traces CG activity in increasing concentrations of glibenclamide (glib.) and after drug wash out. C. Quantification of the number of small cell spikes per burst in different diazoxide concentrations using Friedman-Dunn’s test for repeated measures, post-hoc Wilcoxon-signed rank test with Bonferroni’s correction. Significant difference between-Baseline-200μM: p=0.003; Baseline-400μM: p=0.009. C. Quantification of the number of large cell spikes per burst in different diazoxide concentrations. Same statistical test as C. Significant differences between: Baseline-200μM: p=0.002; Baseline-400μM: p<0.001; Baseline-100μM: p=0.030; 100μM-400μM: p=0.046. D. Quantification of the burst frequency of different glibenclamide concentrations using repeated measures ANOVA ((F,, 1.52)= 23.58; p<0.001), with post-hoc pairwise comparisons and Bonferroni’s correction. Significant differences between-Baseline-50μM: p=0.043; Baseline-100μM: p=0.019; Baseline-200μM: p<0.001; 50μM-100μM: p=0.039; 50μM-200μM: p<0.001; 100μM-200μM: p=0.002. Each dot represents a single preparation connected by lines in different conditions, horizontal bars represent population means. *: p<0.05; **: p<0.01; ***:p<0.001.

### SUR-associated channel genes and gene expression

While the pharmacological evidence of SUR expression in the two networks is compelling, it does not help distinguish between the presence of K-ATP and TRPM4-SUR channels. Differential channel expression could be a potential mechanism for the different outcomes in the two networks. K-ATP channel expression is known to occur constitutively, but TRPM4-SUR channels are only expressed upon injury in mammals^8,24^. Putative sequences for a C. borealis TRPM4 and K_IR_ have previously been identified^25^, which are the two known partners of SURs. Additionally a new putative sequence for SUR was identified from the C. borealis transcriptome using bait sequences from Daphnia pulex and Drosophila melanogaster SUR sequences (GeneBank ID: PP812399). A single SUR sequence was found, and a single isoform(dSUR) is reported in Drosophila as well. The identified sequence bears motifs for the SUR1 transmembrane domains 1 and 2, and nucleotide binding sites. CG and STG tissue samples collected from the same animal were probed for the presence of the mRNA of previously reported sequences of known channel associates and the SUR sequence we identified. SUR, K_IR_ and TRPM4 mRNA expressions are detectable in both the STG and CG at the expected product sizes (138, 135, 194 base pairs (bp) respectively) (Fig 7), and we confirmed this in samples from 4 animals (data not shown). This suggests that either channel could be partnered with SUR in different regions of the C. borealis nervous system and it is not possible to eliminate either candidate on the basis of gene expression alone.

**Figure 7.**
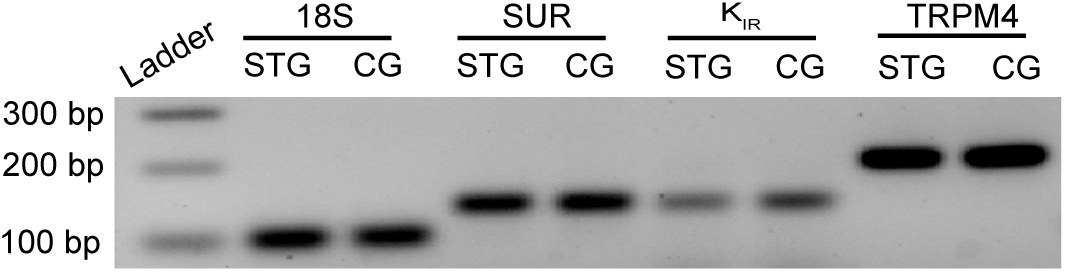
SUR and ion channel expression in the STG and CG: Gel electrophoresis in 2% agarose gel showing amplified PCR products stained with ethidium bromide. Samples were obtained by synthesizing cDNA from extracted mRNA in paired samples of Cancer borealis STG and CG tissue and amplifying the sequences of inquiry via PCR. Lane 1 is a 100bB ladder, Lane 2 and 3 are 18S amplification (reference standard), lane 4 and 5 are SUR, lane 6 and 7 are K_IR_, lane 8 and 9 are TRPM4 PCR amplification products from the same paired STG and CG.

## Discussion

The most striking result in this paper is that pharmacological agents that open or close SUR-associated conductances have diametrically opposing effects in two CPGs in the crab, C. borealis. The pyloric rhythm is inhibited by opening SUR-associated channels whereas the CG’s rhythmic bursting activity increases under the same conditions. Closing SUR-channels in the two networks also has opposing effects, pyloric bursting frequency increases whereas the CG burst frequency decreases. These different outcomes are reminiscent of the effects of K-ATP activation in vertebrate physiology which have very different biophysical properties from the channels described herein.

K-ATP channels open and hyperpolarize hippocampal and striatal neurons preventing all spiking activity^26,27^ when [ATP] decreases dramatically due to hypoxia/ischemia^11,14^, protecting the neurons from excitotoxic death. Pharmacologically opening SUR-conductances in the pyloric network causes the pacemaker kernel neurons to depolarize instead. However, SUR-activation in the STG produces the same outcome as K-ATP opening, all firing and slow wave oscillations stop by the action of a large leak-like shunt. To the best of our knowledge, the STG is the only spontaneously rhythmic neuronal network to be silenced by SUR-activation; many other mammalian networks require K-ATP activation to burst^28,29^. PD neurons can generate slow wave oscillations even when action potentials are blocked^30^; therefore, action potential-related decreases in [ATP]_i_ may not be critical for rhythm generation in this network. Decreased firing when SUR-channels are open may instead be related to STG activity changes in hypoxic challenges experienced during molting in crustaceans, in which the output of the pyloric network reduces with decreasing availability of oxygen^31^.

Closing channels in STGs with intact neuromodulatory inputs produces minor changes in network burst properties. However, the impact of channel closing is much more pronounced in decentralized preparations in which the modulatory inputs are removed. This may be due to ceiling effects in intact conditions, channels could be mostly closed in intact networks due to sufficient [ATP]_i_ levels and more open upon decentralization in response to metabolic shifts. This would imply that decentralization leads to a drop in [ATP]_i_ in pacemaker neurons in the STG which would be an unusual outcome of reduced neuronal activity. The STG is a heavily modulated network, perhaps allowing for a largely invariant output in the face of changing external circumstances^32^. Alternately, removal of these inputs could change the contribution of a single conductance which is otherwise swamped by more prominent neuromodulator-driven conductances which would explain the differences we report.

The crab heartbeat is driven through neural circuitry whereas SUR contributions to cardiac function have been studied primarily through their expression in the myocardium of many animals^33–35^. SUR-activation and inhibition results in activity changes, similar to those elicited in the CG, in other systems in which rhythmic output is crucial to survival such as the pre pre-B1tzinger complex^36^ and murine hearts^37,38^; but it is difficult to know the exact details of how this is achieved in the CG. Unlike the PD neurons, the CG large cells do not depolarize stably in diazoxide unlike PD neurons and instead large, periodic membrane deflections that initiate action potential bursts emerge (personal observations) which stop in TTX. CG neurons have a large number of spikes per burst. Bursts of spikes can cause sufficient ATP decreases to cause K-ATP channels to open^39^; and this recurrent activation can allow switching from tonic to bursting behaviors in other systems^28,40^.

From the gene expression results reported here, it is impossible to predict whether the same channel associations form in the CG and in the STG. The two known channel partners of SURs are expressed in both networks. SURs themselves have orthologues in other arthorpods^41,42^ and SUR pharmacology gratifyingly functions consistently in many animals^43,44^; both channel associations are activated by the same pharmacology. Therefore, we cannot rule out that different channel-SUR pairs are expressed or activated in the two different networks to lead to different physiological outcomes.

The channel biophysics obtained in PD neurons also suggests a completely novel and unusual configuration of SUR-channel association in C.borealis. K-ATP channels conventionally have a reversal potential of K^+^ (∼-80mv) and TRPM4-SUR channels have a reversal potential close to 0mV. In contrast the SUR-associated conductance in PD neurons has a reversal potential close to that of leak conductances. Ion channel expression correlations have been reported to exist widely^45,46^. Therefore, a combination of the two known channel partners in fixed proportions could well explain the channel properties observed. The calcium-sensitivity of the conductance suggests a TRPM4-like channel’s involvement and while the channel is K^+^-permeable, its reversal potential indicates additional ion permeabilities. These properties could also originate from SURs pairing with a TRPM4-like channel^10^ with a reversal potential close to leak, or a completely novel channel pairing yet to be identified. A definitive answer would require molecular protocols that are not currently available for C.borealis. SURs are promiscuous and there could be an unknown channel association that contributes to neuronal biophysics in unexpected ways in different systems.

Our results highlight a dichotomous response from the same receptor/channel activation in different networks. In the crab, it appears that ATP-sensitivity might be required by some networks to produce their desired output and may exist to play a neuroprotective role in others, perhaps allowing the heart to respond in a way that would increase energy availability while simultaneously stopping non-essential stomach pumping to preserve resources. This would be a means of enhancing the physiological resilience of animals to changes in their environment through one common mechanism, although it is also possible that the different effects are an accident of evolution. SUR expression changes in different disease contexts^47,48^ but also in order to facilitate adaptation to extremely hypoxic environments^34^. Oceans are warming rapidly which is concomitant with changes in dissolved oxygen, both of which will impact metabolic rates in all marine life. SUR-dependent mechanisms could provide a host of protective mechanisms that can help crabs adapt and acclimate to multiple and simultaneous fluctuations in their environment.

## Acknowledgements

We thank Abigail Zuber for her help with primer development and early PCR assays, Maria Ivanova and Naamah Romano for help with electrophysiology data collection, Nevo Schpire for pilot experiments, and Katelyn Wadland for dissection assistance.

The authors report no conflicts of interest

Funding source: R35 NS097343

## Notes

### Competing Interest Statement

The authors have declared no competing interest.

